# EECoG-Comp: An Open Source Platform for Concurrent EEG/ECoG Comparisons: applications to connectivity studies

**DOI:** 10.1101/350199

**Authors:** Qing Wang, Pedro Antonio Valdés-Hernández, Deirel Paz-Linares, Jorge Bosch-Bayard, Naoya Oosugi, Misako Komatsu, Naotaka Fujii, Pedro Antonio Valdés-Sosa

**Affiliations:** The Clinical Hospital of Chengdu Brain Science Institute, MOE Key Lab for Neuroinformation, University of Electronic Science and Technology of China, Chengdu, China; Department of Biomedical Engineering, Florida International University, Miami, FL, United States; Unity of Neurodevelopment, Institute of Neurobiology, UNAM, Campus Juriquilla, Santiago de Querétaro, Querétaro, México; Laboratory for Adaptive Intelligence, RIKEN Brain Science Institute, Saitama, Japan; Laboratory for Molecular Analysis of Higher Brain Function, RIKEN Center for Brain Science, Saitama, Japan; Cuban Neuro Science Center, La Habana, Cuba

**Keywords:** EEG, ECoG, Forward Modeling, Electrophysiological Source Imaging, Connectivity, Open Science, Monkey electrophysiology

## Abstract

Electrophysiological Source Imaging (ESI) is hampered by lack of “gold standards” for model validation. Concurrent electroencephalography (EEG) and electrocorticography (ECoG) experiments (EECoG) are useful for this purpose, especially primate models due to their flexibility and translational value for human research. Unfortunately, there is only one EECoG experiments in the public domain that we know of: the Multidimensional Recording (MDR) is based on a single monkey (www.neurotycho.org). The mining of this type of data is hindered by lack of specialized procedures to deal with: 1) Severe EECoG artifacts due to the experimental produces; 2) Sophisticated forward models that account for surgery induced skull defects and implanted ECoG electrode strips; 3) Reliable statistical procedures to estimate and compare source connectivity (partial correlation). We provide solutions to the processing issues just mentioned with EECoG-Comp: an open source platform (https://github.com/Vincent-wq/EECoG-Comp). EECoG lead fields calculated with FEM (Simbio) for MDR data are also provided and were used in other papers of this special issue. As a use case with the MDR, we show: 1) For real MDR data, 4 popular ESI methods (MNE, LCMV, eLORETA and SSBL) showed significant but moderate concordance with a usual standard, the ECoG Laplacian (standard partial *AUC* = 0.65 ± 0.05); 2) In both monkey and human simulations, all ESI methods as well as Laplacian had a significant but poor correspondence with the true source connectivity. These preliminary results may stimulate the development of improved ESI connectivity estimators but require the availability of more EECoG data sets to obtain neurobiologically valid inferences.

## 1. Introduction

Among the non-invasive technologies used to probe normal and pathological brain function in both humans and animals, Electroencephalography (EEG) (Schomer and Da Silva 2012) is the most affordable, popular and widespread. It also provides the highest temporal resolution. Alas, like any other imaging modality, EEG is not a straightforward proxy for neural activity. In fact, it is a spatially filtered and sub-sampled reflection of hidden brain states, corrupted as well by noise and artifacts. Indeed, at the EEG electrodes, the signals are spatially filtered by volume conduction (van den Broek et al. 1998)—owing to the very low conductivity of the skull, and they are the observation of mixed EEG source activities. This mixing also introduces spurious brain coherence, leading to false brain connectivity estimations. For this reason, EEG connectivity should be, in principle, assessed from estimates of source activity. Thus, what is required is the solution of the inverse problem (Pascual-Marqui 1999), i.e. the estimation of brain sources activities from EEG recordings—a technique also called Electrophysiological Source Imaging (ESI).

ESI has been long studied in the attempt to localize epileptic foci (Ding et al. 2007; He et al. 2011). In addition, it has been proposed to study the localization of different brain functions as well as the dynamics of neural connectivity. Nevertheless, ESI methods have important methodological issues and requires external validation (Nunez et al. 2019).

Electrocorticography (ECoG) or intracranial electroencephalography (iEEG) is considered by many as a “gold standard” in clinical settings, since it provides more accurate information of the cortical brain states (Nunez et al. 2019; Frauscher et al. 2018). ECoG is a type of invasive monitoring technology with electrodes placed directly on the exposed surface of the cortex to record electrical activity of the brain. It should be noted that, even though the ECoG electrodes are directly placed on the cortex, the activity at the surface of the cortex is not equivalent to the true “sources” of the “physiological cortical macro columns” proposed by Buxhoeveden et al. (2002). This is the reason why we need to solve the inverse problem for ECoG as well (Nunez et al. 2019).

Solving the inverse problem first requires the solution of the forward problem (Hallez et al. 2007) for EEG and ECoG (EECoG), i.e., the calculation of the so-called lead field matrix (LF), a mapping matrix from known sources to EECoG measurements. Such mappings demand the definition of a head model specifying the spatial distribution of tissue conductivity, a source space model and the location the EECoG electrodes with respect to the source space. The more realistic the head model, the more accurate the ESI solutions. Nowadays, high detailed structural Magnetic Resonance Images (MRI) are used to define the head model and either Boundary Element Method (BEM) or Finite Element Method (FEM) are used to calculate the LF. These methods differ in accuracy.

The EECoG inverse problem is ill-posed in the Hadamard sense: the null space of the lead field has finite dimension, sabotaging the uniqueness of the solution. Thus, it is necessary to impose priors on the sources such as minimum energy, sparsity, smoothness and independence; and to restrict the source space with anatomical information, e.g. by forcing the sources to be in the gray matter or by imposing the orientation of the dipoles to be perpendicular to the cortical layer. This is another reason to provide accurate forward head models. Abundant inverse solutions using of these constraints have been developed and implemented, examples being weighted minimum-norm estimation (WMNE) (Hämäläinen et al. 1994), beamforming (BF) (Grech et al. 2008; Van Veen et al. 1997), Blind Source Separation (BSS) (Oosugi et al. 2017), exact Low Resolution Electromagnetic Tomography (eLORETA) (Pascual-Marqui 2007), Sparse Structural Bayesian Learning (SSBL) (Paz-Linares et al. 2017), and so on. This problem is even more difficult for source connectivity estimation, as an aspect that we will later address more closely.

It follows, due to the complexity of the forward modeling procedure and the diversity of possible inverse solutions, we can arrive at completely different source activity and connectivity estimations from different ESI methods. In view of the proliferation of ESI methods, it is quite challenging to determine which is the best and to quantify its properties reliably. There is general agreement that simulations are not enough to make this decision—experimental verification is necessary.

Unfortunately, there is a paucity of data sets that can serve as a “gold standard” to evaluate ESI. This is unsurprising given the obvious difficulty in recording concurrent scalp EEG signals in a normal intact head simultaneously with direct brain activity measurements. This problem was only tackled with a few unrealistic (though exceptionally useful) attempts in the past. That is the case of Leahy et al. (1998) who designed and conducted a multiple dipole phantom consisting of 32 independently programmable and isolated dipoles inserted in a skull mount filled with conductive gelatin. They used this phantom to collect EEG and MEG data to evaluate the performance of current dipole localization methods. In Marin et al. (1998) another realistic conductive skull phantom was used to evaluate the accuracy of forward FEM methods. Later, this phantom was used to evaluate the performance of different ESI methods in Marin et al. (2001)^2^. Recently, Peterson and Ferris (2018) evaluated the performance of ICA-based connectivity in a phantom in motion. A much more ideal “phantom” to study the effect of the accuracy of ESI methods on assessing brain connectivity and source localization is the one providing scalp EEG signals concurrently with actual and direct whole brain measures of brain activity, such as Local Field Potential (LFP) and Multi-Unit Activity (MUA) (Mattia et al. 2010). However, coverage of all brain areas simultaneously with these techniques is impossible. On the other hand, optical imaging (Wang et al. 2003) provides a direct measurement of hemodynamic responses at the cortical level, but all these techniques require the removal of the skull during the experiment.

A better candidate is a simultaneous EECoG experiment which allows recording electrical activity directly on the cortex and obtaning simultaneously the scalp EEG after closing the skull. This can achieve comparatively large coverage for both types of recordings. However, the invasiveness makes it available only in human clinical and preclinical settings, mainly restricted to pathologies like epilepsy—only targeting the seizure onset zones and thus usually precluding the recording of larger spatial extensions of the cortex (Frauscher et al. 2018).

A better situation is achievable in primate preparations. For example, the Multidimensional Recording (MDR) database (http://neurotycho.org) (Nagasaka et al. 2011) offers such a kind of simultaneous EECoG data, with ECoG covering the left hemisphere of the cortex and EEG covering the whole head. To our knowledge, this is the only open macaque simultaneous EECoG data set currently available. Despite the complexity of setting up this type of experiment, EECoG has the unique advantage of allowing cross validation of two types of recordings originated from the same neural sources. Testing the ESI methods has also been possible with this dataset (Oosugi et al. 2017). Even though the electrophysiological and imaging data is limited to a single monkey, it is the only publicly available dataset with the required information for detailed forward EEG and ECoG modeling.

Nevertheless, there is not yet a Neuroinformatics platform (pipeline) designed for analyzing and comparing such kind of simultaneous EECoG data, especially for the evaluation of source connectivity estimation. The main reasons are:

1. The artifacts induced by the very complicated primate experimental configurations, which hinders the correct estimation of brain sources activity and connectivity patterns;
2. The difficulty in obtaining a realistic head model that takes into consideration the implanting of very low conductivity ECoG silicone stripes, and the difficulty of computing the lead fields for EEG and ECoG with the same source space;
3. The lack of tools to compare results in terms of connectivity estimation with the same level of statistical significance.

In this paper, we solve these problems and provide a public and open source platform named “EECoG-Comp” that has the following features:

1. Customized preprocessing algorithms, including synchronization and model-based artifact removal algorithms;
2. Realistic MRI-based biophysical head model which takes into consideration of the implanted ECoG stripes, and provides FEM-based lead fields for both EEG and ECoG with the same source space;
3. Standard source connectivity simulation and statistical analysis to guarantee the reliability of comparisons.

This platform is shared via https://github.com/Vincent-wq/EECoG-Comp, and the data will be available upon request. For a detailed description of the content shared, refer to Table 2 in next section. The lead fields from our platform have been used to research directed connectivity measures (Papadopoulou et al. 2015) and for testing the resolution properties of different inverse solutions (Todaro et al. 2018).

**Table 1.** The inverse methods implemented in ESI submodule.

| Acronym | Name | Reference |
| --- | --- | --- |
| MNE | Minimum Norm Estimation | Hämäläinen et al. (1994) |
| LCMV | Linearly Constrained Minimum Variance Spatial Filtering | Van Veen et al. (1997) |
| eLORETA | Exact Low Resolution Electromagnetic Tomography | Pascual-Marqui (2007) |
| SSBL | Sparse Structural Bayesian Learning | Paz-Linares et al. (2017) |

**Table 2.**
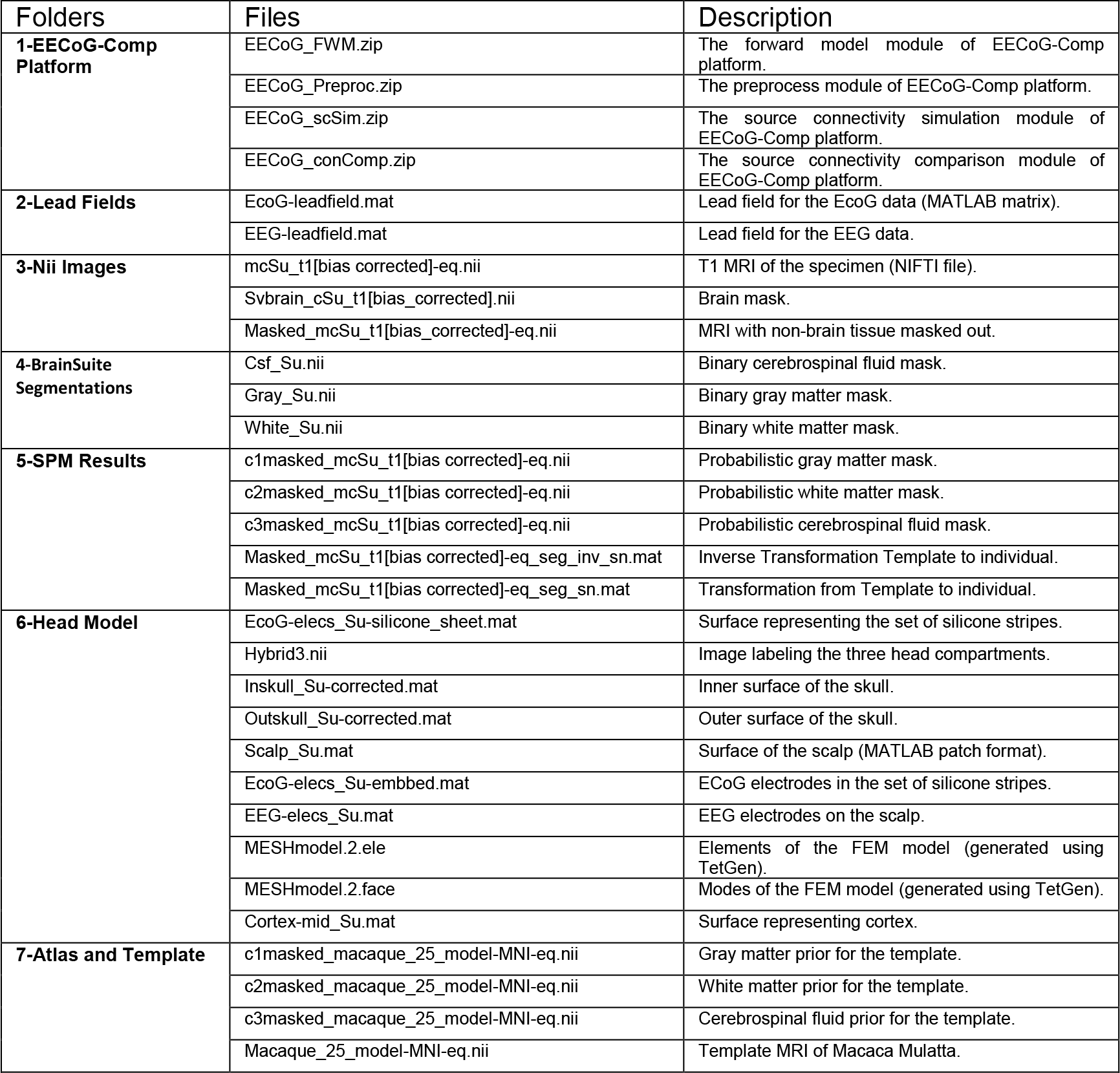
Table of shared contents from EECoG-Comp platform.

In this paper, we describe the simultaneous EECoG platform in section 2. As an illustration of the usefulness of this platform, we take ESI methods connectivity comparison as an example in section 3 and 4. Our preliminary results yield interesting results, indicating that the performance of the tested ESI methods seem to detect connectivity slightly above chance but with very low accuracy.

## 2. Concurrent EEG/ECoG Comparison Open Source Platform

The concurrent EEG/ECoG comparison open source platform (**EECoG-Comp**) is composed of four main modules as depicted in Fig. 1, and they are: 1) **EECoG_Preproc** (Fig. 1 upper right) for preprocessing; 2) **EECoG_FWM** (Fig. 1 lower left) for realistic forward modeling; 3) **EECoG_scSim** (Fig. 1 upper left) for EECoG source connectivity simulation; 4) **EECoG_conComp** (Fig. 1 lower right) for inverse solution, graph model estimation and statistical comparison. They are designed respectively for: 1) Rejecting the artifacts induced by the very complicated experimental configurations; 2) Solving the EEG forward problem in presence of the very thin silicone ECoG subdural array; 3) Create simulations as null model for testing of the methods (for example ESI methods); 4) Solving the inverse problem, estimating and comparing the source level connectivity of EECoG at the same significance level.

**Figure 1.**
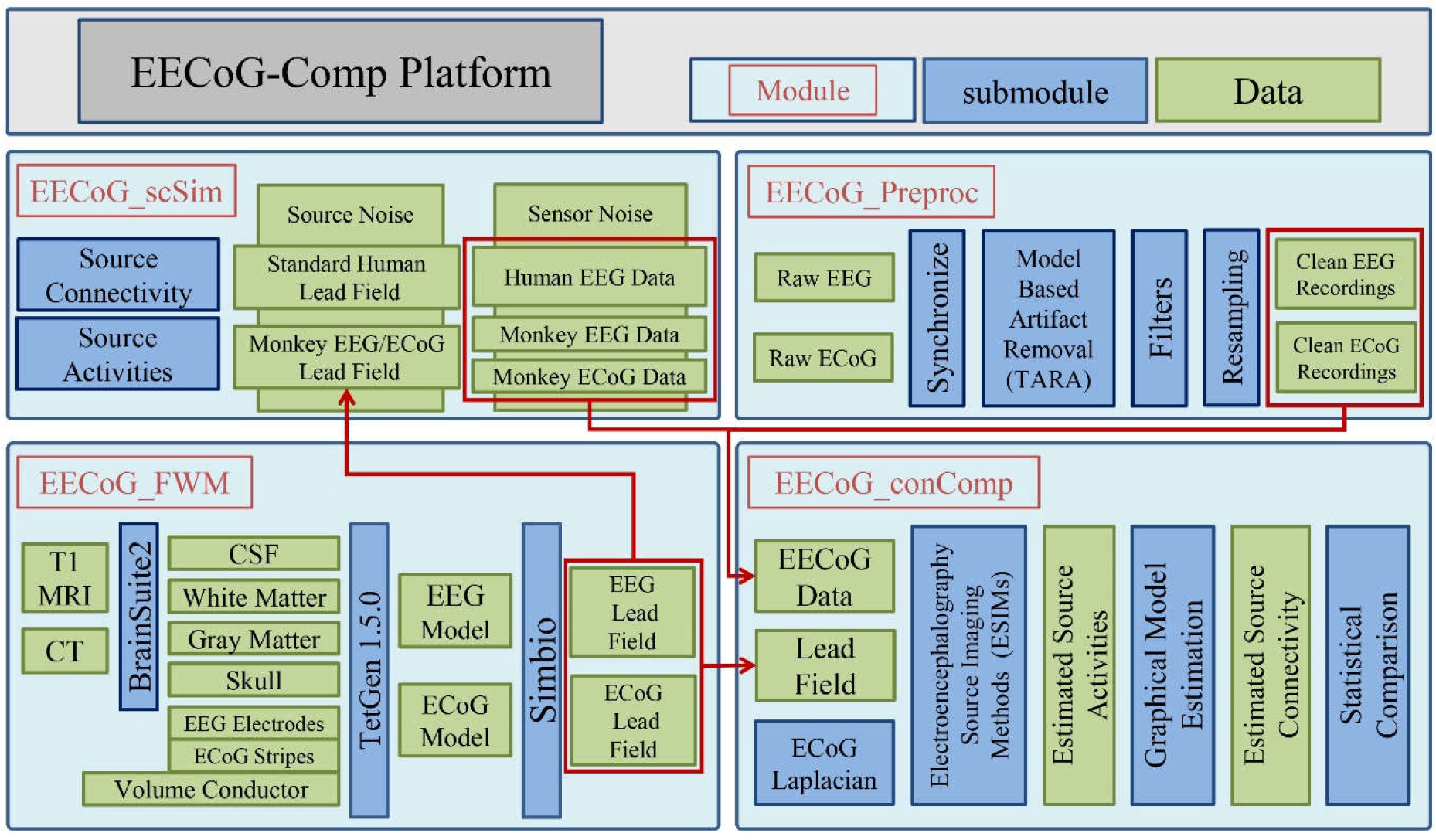
System diagram for the open source EECoG-Comp platform.

In what follows, we describe the modules and submodules relevant to specific challenges of EECoG connectivity analysis. We take the simultaneous EECoG experiment from neurotycho.org (http://neurotycho.org/eeg-and-ecog-simultaneous-recording) as an example to illustrate how EECoG-Comp works. This is shown in the following subsections. The following mathematical notations are used to explain the theory and methods behind all these modules: *x* ∈ ℝ (lower case, Italian, not bold) is a function or a variable, ***x*** ∈ℝ^*N*^ (lower case, Italian, and bold) is a vector, ***X*** ∈ ℝ^*M*×*N*^ (upper case, Italian, and bold) is a matrix or matrix operator, refer to Table 3 in the Appendix 1 for all details of the notations.

### 1 MDR Recording

This dataset is partly described in Nagasaka et al. (2011) and Oosugi et al. (2017). All the experimental and surgical procedures were performed in accordance with protocols approved by the RIKEN ethical committee (No. H22-2-202(4)) and the recommendations of the Weatherall report, “the use of non-human primates in research”. During the experiment, a macaque monkey (*macaca fuscata*) was seated in a primate chair with hands tied and head movement restricted. The eyes of the monkey were covered to avoid evoked visual responses during the entire experimental period.

The electrophysiological data (EEG and ECoG) was recorded during two conditions: 1) awake resting state with eyes folded and movements restricted; and 2) drug induced anesthesia: 1.15 ml (8.8 mg/Kg) ketamine hydrochloride and 0.35 ml (0.05 mg/Kg) medetomidine injection. Two five-minute trials of simultaneous resting state EEG and ECoG activities were recorded for each condition.

For ECoG, the skin, skull and dura of the monkey’s left hemisphere was carefully removed. Then, a modified human subdural ECoG array, adapted to the monkey physiology (Nagasaka et al. 2011), was inserted in the subdural space between the arachnoid membrane and the dura mater (see Fig. 2). This array was fixed on a set of 1 mm-thick silicone stripes, containing 128 insulated ECoG platinum disk-shaped electrodes with 3 mm in diameter and leaving only a hole in the silicone of 0.8 mm diameter at the center of each electrode. The electrodes, with a separation of 5 mm between each other, covered the frontal, parietal, temporal and occipital lobes and part of the medial wall of the left cortex, as depicted in the X-ray image in Fig. 3. The X-ray images were obtained from a VPX-30E system (Toshiba Medical Supply, Tokyo, Japan) with 8.0 mA in current and 90 kV in voltage. The reference electrode were rectangular platinum plates in the subdural space between the array and the dura. Another platinum electrode was placed as ground in the epidural space.

**Figure 2.**
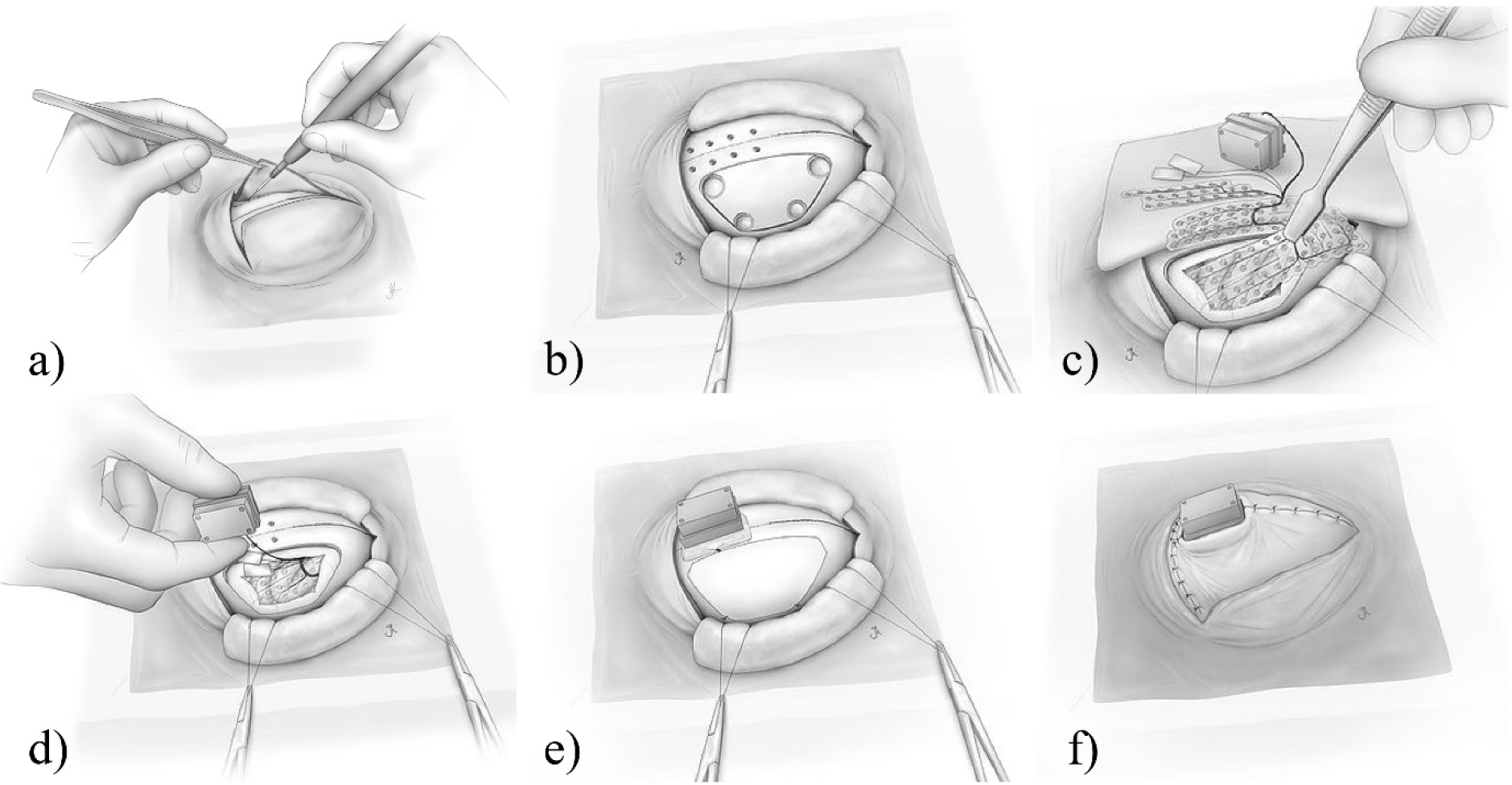
Illustration of the procedure for implanting ECoG arrays and placing back the dura, the skull and the skin to cover the brain cortical external surface.

**Figure 3.**
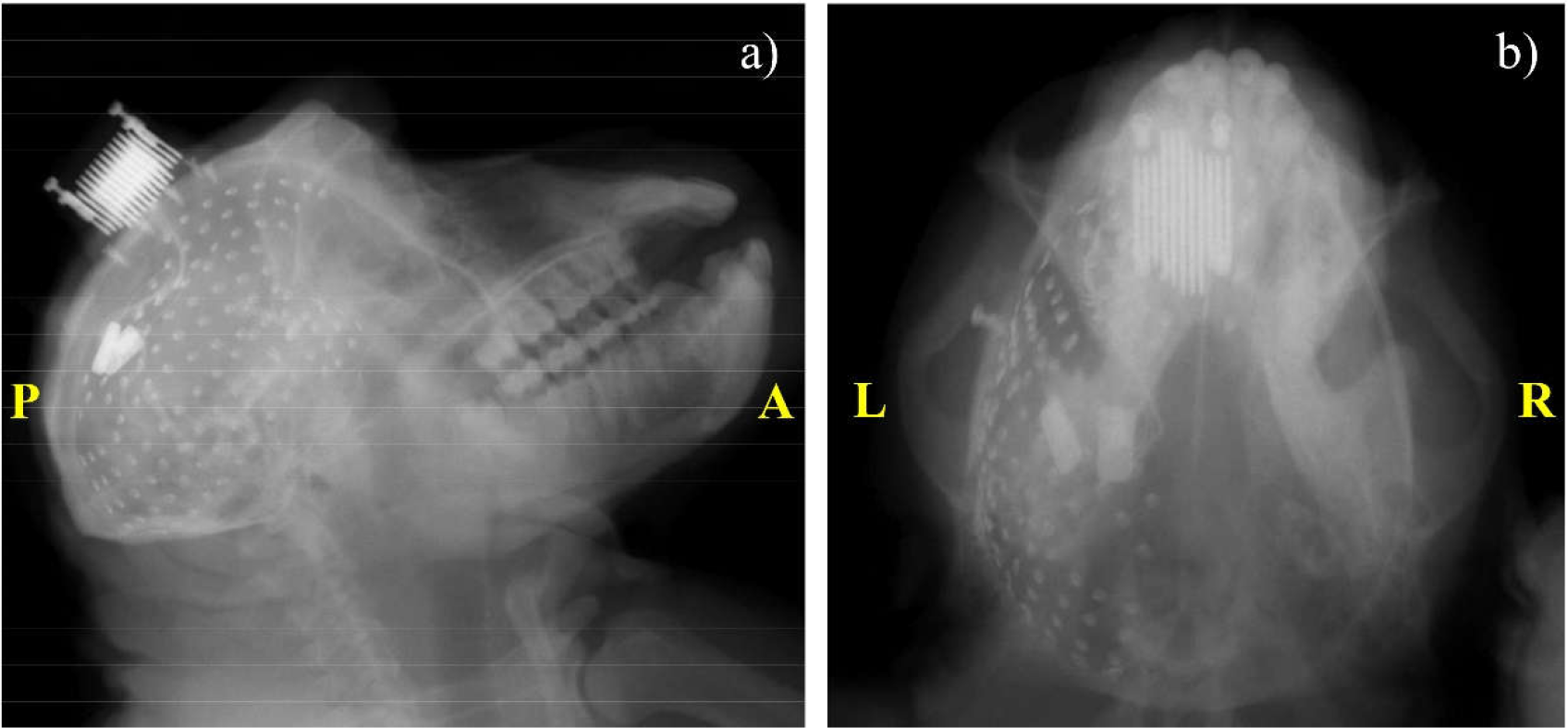
X-ray images of the ECoG electrodes: 1) Sagittal view; b) Coronal view.

After placing the ECoG electrodes, the dura, the skull and the skin were carefully placed back (see Fig. 2). ECoG was filtered and recorded by a Cerebus recording system (Blackrock Microsystems, Utah, USA). Plastic connectors from the array to the ECoG system were attached to the top of the head with dental cement and small titanium screws. The sampling rate was 1000 Hz, and the recordings were Butterworth bandpass filtered within 0.3-400 Hz.

The EEG was recorded with a NeuroPRAX system (eldith GmbH, Ilmenau, Germany) of sampling rate 4096 Hz, using 18 electrodes which follows the standard 19-electrode 10-20 system. Electrode “Cz” has been removed to place the ECoG plastic connector. The recordings were down-sampled to 1000 Hz. An external TTL pulse was used to define the beginning and the end of the EEG and ECoG recordings interval of interest for synchronization.

### 2 Preprocessing module: EECoG_Preproc

The first challenge of analyzing the EECoG (MDR) data is the high level of artifacts present in the recordings. We therefore addressed this attention to assimilating procedures to alleviate this interference.

The first preprocessing step of the simultaneous EECoG recordings is synchronization. This is done by aligning the original 4096 Hz EEG and 1000 Hz ECoG recordings to the “Trigger” signal. After that, the “Trigger” channels are removed from both recordings. Then, we average referenced the recordings and remove the mean. Based on the observed EEG artifact properties of sparse spikes and step-like discontinuities, we further use Transient Artifact Reduction Algorithm (TARA, the theory will be detailed in Appendix A2) (Selesnick et al. 2014) to clean these artifacts, the results are shown in Fig. 4 below. We further apply the notch filter to remove the power-line artifact of 50 Hz and applied the Butterworth high-pass filter of 0.3 Hz. At last, we down-sampled the EECoG recordings to 400 Hz and bandpass to alpha band (8 Hz to 12 Hz) for latter analysis. Our platform also supports FieldTrip (Oostenveld et al. 2011) and EEGLab (Delorme and Scott 2004) data structure, which allows the users to create their own pipeline out of this platform.

**Figure 4.**
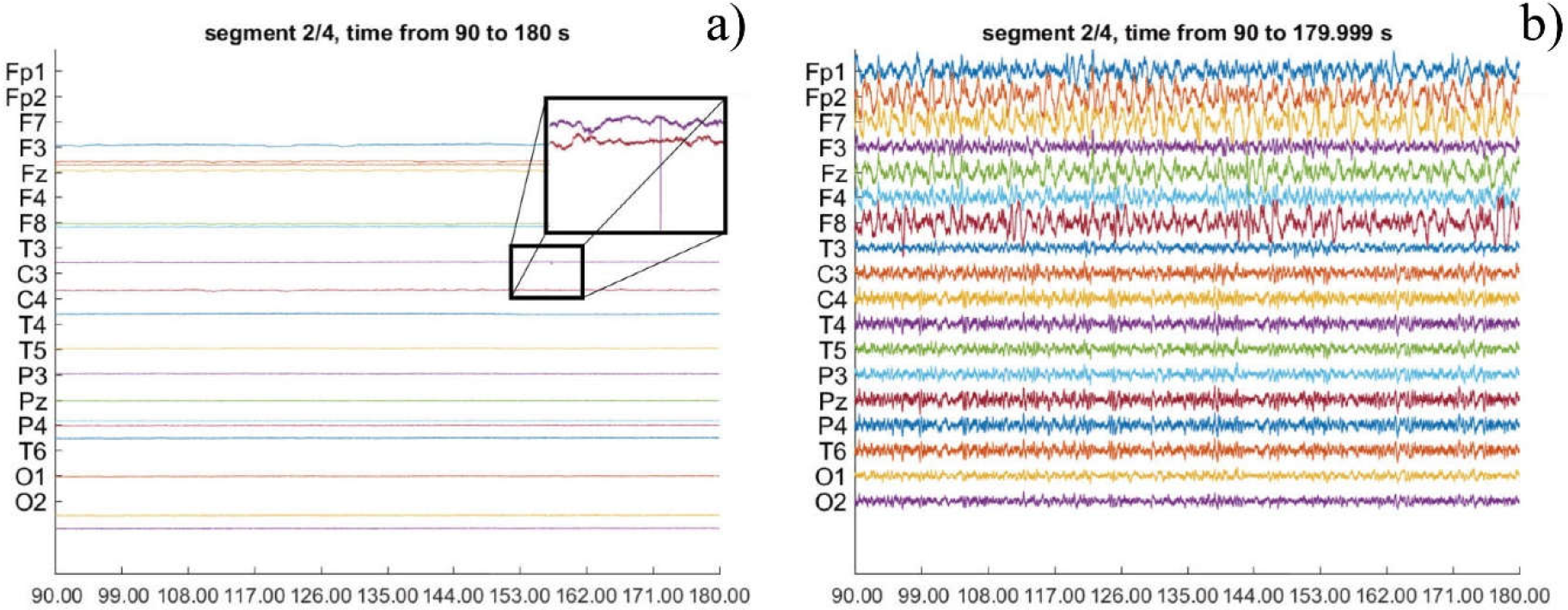
The comparison of the EEG recordings before and after preprocessing with TARA. (a) The original EEG recordings of anesthesia trial 5 (Y scale is [-2909.6, 2690.4] unit), the zoomed in window shows the artifacts of sparse spikes and step-like discontinuities in the original recordings. (b) The cleaned EEG recordings of anesthesia trial 5 (Y scale is [-60, 60] unit).

By introducing TARA, we were able to clean most of the above two types of artifacts as shown in Fig. 4. After preprocessing, several channels in the EEG (F3, C3, and Fz) and ECoG (channel 38, 50, and 93) have been removed from further analyzing due to bad quality, as visually judged by an expert neurophysiologist.

It is to be noted that, surprisingly, we did not observe a clear alpha peak, not even from the occipital recordings of both EEG and ECoG, despite our careful artifact rejection and pre-processing.

### 3 Forward Modeling Module: EECoG_FWM

The second challenge of simultaneous EECoG data analysis comes from the implanted ECoG stripes. This makes the forward modeling procedure much more difficult. EEG forward modeling is the first step for Electrophysiological Source Imaging (ESI) analysis, and it can provide the so-called lead field or gain matrix ^***K***^ for further source activity estimation and simulation. Thus, EECoG_FWM, as one important module of EECoG-Comp, is created to build the realistic head model for the complex simultaneous EECoG experiment. The EEG forward problem can be formulated as follows:

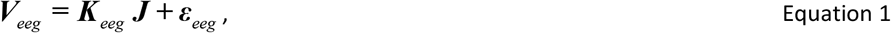

While the ECoG forward problem can be written as:

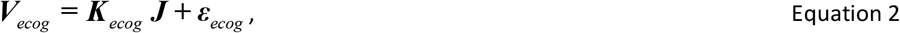

where ***J*** is the primary current density, ***V**_eeg_* and ***V**_ecog_* are the measured EEG and ECoG signals, ***K**_eeg_* and ***K**_ecog_* are the lead fields for EEG and ECoG, ***ε**_eeg_* and ***ε**_ecog_* are the measurement noises for EEG and ECoG, respectively.

The aim of EECoG_FWM module is to estimate ***K**_eeg_* and ***K**_ecog_* from the T1-weigthed Magnetic Resonance Image (MRI) of the subject (in this data set, it is the only monkey named Su), these images were used to construct the surfaces (triangular meshes) defining the head model. The T1w image was acquired in a 4T Varian Unity Inova MR scanner (Varian NMR Instruments, Palo Alto, CA) with a 3 in 1 loop surface coil (Takashima Seisakusho Ltd., Tokyo, Japan). The scanning parameters were: TR/TE = 13 ms/3.8 ms, in-plane matrix of 256 × 256 × 256, field of view (FOV) of 12.8 × 12.8 × 12.8 *cm* with voxel size of 0.5 × 0.5 × 0.5 *mm*. This image was segmented into white matter (WM), gray matter (GM) and cerebrospinal fluid (CSF) with BrainSuite2 (Shattuck et al. 2001). The gray and white matter interface surface, as shown in Fig. 5(a), is chosen as the source space model for EECoG, i.e. each of its 104650 nodes can be the location of a current dipole with orientation normal to the surface, and we are using a down-sampled version in latter experiments due to the computational issues. The volume conductor model comprises three compartments 1) WM+GM+CSF, 2) Skull and 3) Skin as shown in Fig. 5(a) with constant conductivities 0.33 S/m, 0.0041 S/m and 0.33 S/m, respectively. These are represented by their boundaries: the “inskull”, the “outskull” and the “scalp”. Additionally, a fourth compartment accounting for the ECoG silicone stripes were included, with 1 mm thickness and the shape shown in Fig. 5(b), and they are also described in the first subsection named “MDR Recordings”. It is very important to account for this compartment because its low conductivity and therefore it influences the EEG signals. On the other hand, the EEG electrodes were manually located on the monkey’s scalp using IMAGIC software (www.neuronicsa.com) as depicted in Fig. 5(c).

**Figure 5.**
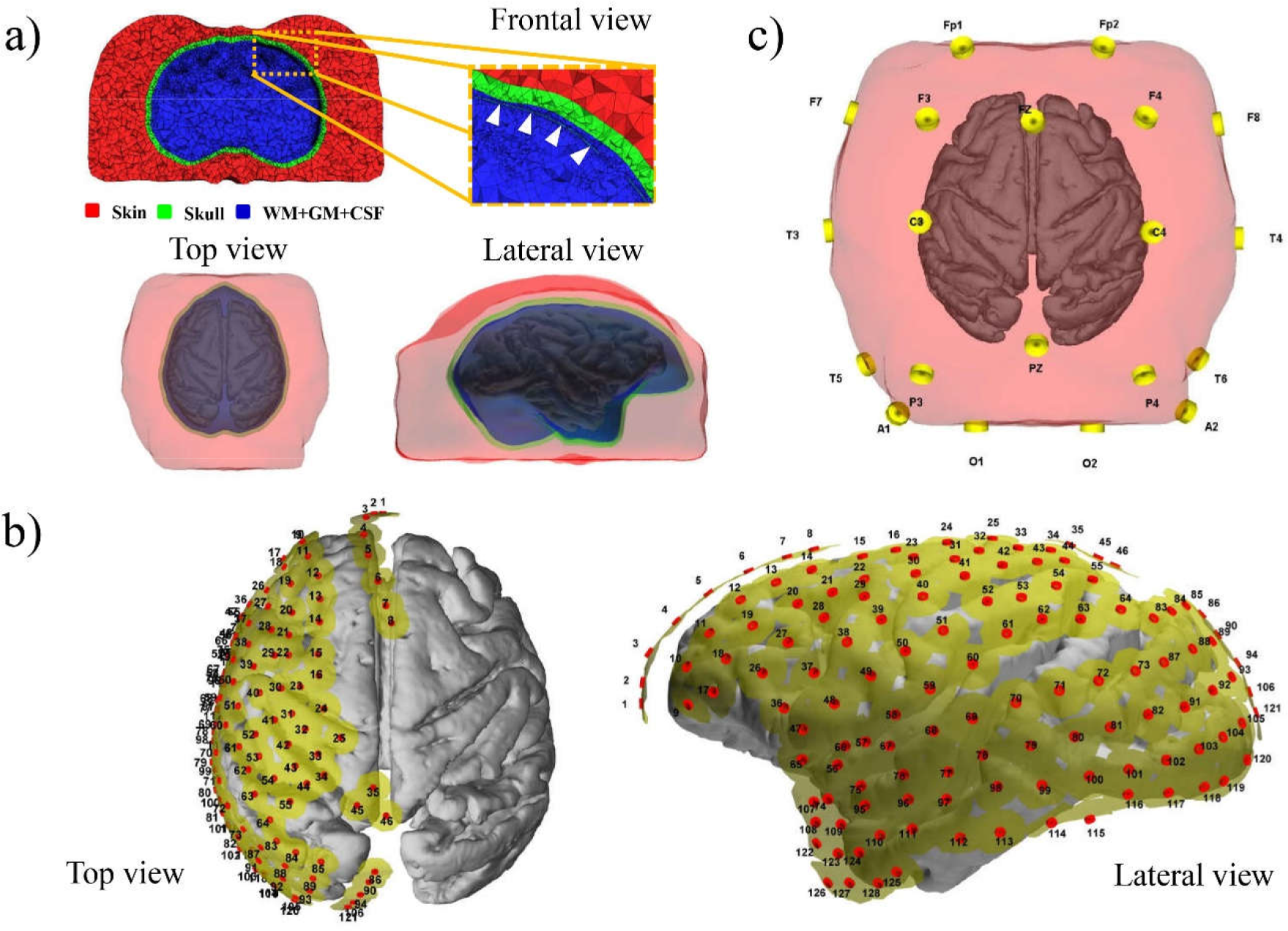
Head Model of the monkey: a) Tetrahedral meshes defining the source space and the volume conductor model. From inside to outside: source space (cortex), inskull, outskull, and scalp; b) silicone compartment (yellow) and 128 ECoG electrode positions represented (source space is shown as reference); c) The 10-20 system EEG electrodes with the head model.

The tetrahedral meshes in Fig. 5(a) were created from the surfaces of the head model using TetGen 1.5.0 (Si 2015), an open source software in Linux repositories. During the meshing process the nodes in the source space surface (cortical grid) were forced to be nodes of the tetrahedral mesh to guarantee the Venant’s condition. Both EEG and ECoG lead fields were calculated using NeuroFEM. This is a program for computing lead fields using FEM which is part of the SimBio software package (Consortium et al. 2000) (index page: https://www.mrt.uni-jena.de/simbio/index.php/Main\_Page). This module can provide average referenced LFs based on 3d cortical grid source space.

It should be noted that only 15 channels of EEG left after removing the bad channels, and 125 channels of ECoG left. Finally, we only took the left hemisphere of the cortical grid (source space) and study its connectivity due to the limited coverage of ECoG in real data analysis.

### 4 Source Connectivity Comparison Module: EECoG_conComp

Brain connectivity has received much attention in recent years (Bassett DS et al. 2017). However, the estimation and comparison of brain connectivity at source level is still very challenging (Zalesky et al. 2010). The next challenge for EECoG-Comp platform is to compare the source level connectivity with sound statistics. EECoG_conComp is designed to fulfill this goal, and it has the following submodules: 1) ESI methods submodule, which provide the unified interface for all the ESI methods to estimate the source activities; 2) Graphical model estimation submodule, which estimate the graphical model from the estimated source activities; and 3) The statistical comparison submodule, which compare the estimated graph model parameters at the same statistical significance level. In addition to all these submodules, EECoG_conComp also provides the ECoG Laplacian submodule, which estimates the Laplacian of ECoG over the cortical grid as another way of exploring the “pseudo source connectivity”. All these submodules will be detailed below.

#### ECoG Laplacian Submodule

With its wide acceptance, we shall use the ECoG Laplacian over the cortical grid as a surrogate measure for the primary current density ***J***. Note that, this estimator is very similar to the “spline Laplacian/dura images” discussed by Nunez and Srinivasan (2006).

The specific challenge of this submodule is to approximate the Laplacian over the same cortical grid that will be used for ESI methods. During this procedure, we are assuming that the ECoG Laplacian is smooth over the cortex. Suppose that ***V**_cortex_* is the true voltage on the cortex for our cortical mesh, and the Laplacian on the cortex is *ϕ*_c_. Thus, we have the voltage on the cortex (***V**_cortex_* = ***L***^−1^ ⋅*ϕ*_c_), where ^***L***^ is the Laplacian operator, and ***L*** ^−1^ is the inverse Laplacian operator. Then the signals measured by ECoG are written as:

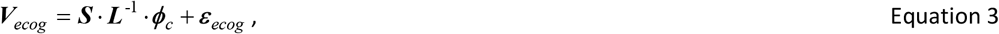

Where ***S*** is a selection matrix that picks the voltage on the full cortical grid which are closest to the ECoG electrodes, and *ε_ecog_* is the ECoG sensor noise with multi-variate Gaussian I.I.D. distribution. We emphasis that the ECoG Laplacian does not depend at all on the knowledge of the realistic head model or lead field. However, ***K**_Lap_* = ***S*** ⋅ ***L***^−1^ can be considered as a type of “pseudo lead field”, and analogues to most inverse problems, Eq. 3 is ill posed, and in order to estimate the Laplacian *ϕ*_c_, we formulate this problem by penalizing the smoothness of the cortex Laplacian, then the problem can be solved by minimizing the following function:

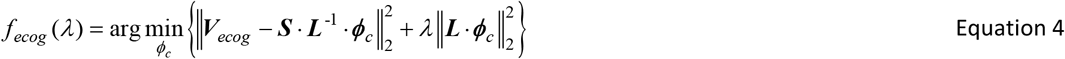

This is a ridge type regression problem, which minimize the estimations square error with penalization of the 4th order Laplacian of the cortex voltage, or the 2nd order Laplacian on the cortex. In order to solve the problem defined in Eq. 3 and Eq. 4, we substitute with ***X*** = ***S*** ⋅ ***L***^−2^ and *Φ*= ***L*** ⋅*ϕ*_c_. Then, Eq. 4 can be rewritten in the standard ridge regression form as:

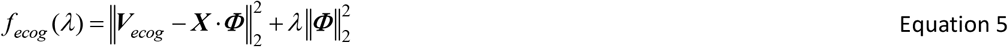

After, minimizing this function and changing variables back, we obtain the estimator for the ECoG Laplacian on the cortical grid:

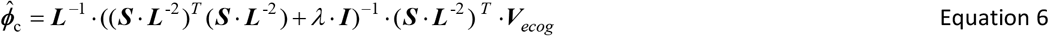

The hyper parameter *λ* is selected with generalized cross validation (GCV). The ECoG Laplacian estimator is calculated for each point of the cortical grid at each time instant, thus also yielding a “cortical grid time series” like the ESI method.

#### Electrophysiological Source Imaging Methods Submodule

This submodule provides a uniform interface to the solution of the EECoG inverse problems and the estimators of the source current ***J***. At present, the inverse solutions implemented are listed in Table 1. These methods completely specified in the references given, and the METH toolbox provides the implementations as well. (https://www.uke.de/english/departments-institutes/institutes/neurophysiology-and-pathophysiology/research/working-groups/index.html).

To compare the source connectivity, we need to ensure that all ESI methods reached their best performance. For MNE, LCMV and eLORETA, we use GCV to select the hyper parameters. For SSBL, the best performance is guaranteed by selecting the hyper parameters with its marginal likelihood stopping criteria.

Each type of inverse solution is calculated for each point on the cortical grid at each time instant. As mentioned before, we need to solve the inverse problem for both EEG and ECoG, then we can compare them on the same source space.

#### Connectivity Estimation Submodules

There are many proposed measures for brain connectivity, both directed and nondirected. We limit our attention of this platform to partial correlations as an undirected brain connectivity measure. This measure has been shown to be less sensitive to indirect connections (Dawson et al. 2016). They can be easily obtained from the inverse of the covariance matrix or precision matrix, which can be normalized to produce the partial correlation matrix (PCM). There are two specific challenges to be solved for this module:

1. The high dimensionality of the covariance matrices to be studied;
2. The lack of standardized statistical criteria for the selections of significant partial correlations.

We here take advantage of recently developed theory for inference of high dimensional sparse precision matrices from Jankova et al. (2018) to address these challenges. The assumption of sparsity seems to be natural and widely adopted for estimators of brain connectivity and is relaxed with recent “debiasing” techniques explained below.

The input for the connectivity estimation submodule is the estimated time series at of the cortical grid from either ESI methods or the ECoG Laplacian. These time series are filtered to a specific frequency band of interest, like alpha of 8-12Hz. From these filtered time series, we calculate the empirical covariance matrix. This empirical covariance matrix is then standardized to produce the empirical correlation matrix 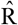. Then, both are used to obtain a sparse estimate of the partial correlations, also known as the weighted inverse covariance 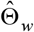 which is estimated by the following minimizing function:

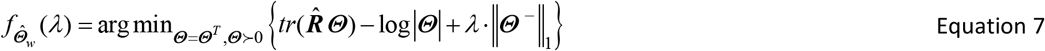

where ||Θ^−^||_1_ is the ***l**_1_* norm of the off-diagonal elements of the inverse covariance matrix *Θ*, and *λ* is the hyper parameter. The computation is readily carried out by the QUIC algorithm (Hsieh et al. 2014), which is one optimized implementation of the graphical LASSO.

We leverage the recent theoretical results from Jankova et al. (2018) in two ways:

First, we can correct the bias introduced by the graphical LASSO with the “debiasing” operation:

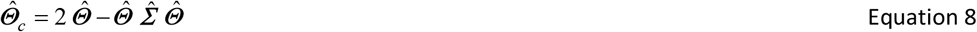

Second, asymptotic normality of the inverse covariances estimator is achieved as stated in Theorem 3 of Jankova’s paper (Jankova et al. 2018) with the following transformation:

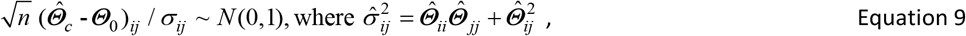

This result allows us to find the thresholds for all the PCMs with the same statistical significance level.

Since we are dealing with very high dimensional PCMs, the threshold must be corrected for multiple comparisons, and it is done by the application of the positive False Discovery Rate *pDFR* (Storey 2002) as follows:

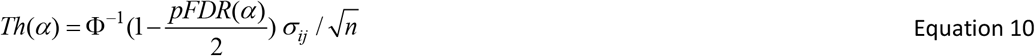

#### Connectivity Comparison Submodules

The connectivity comparison submodule offers mainly 2 methods to compares the thresholded estimated PCMs against a “base connectivity” matrix (either the truth known from simulations or any of the ESI based source connectivity estimators).

The first measure is based on ROC curves, which shows the true positive rate (TPR) against false positive rate (FPR) at all possible FPR values. In order to evaluate the significance of ROC curve, we also calculate the pointwise confidence bound for all the possible FPRs by bootstrap. We correct the bias introduced from the bootstrap distribution with bias-corrected and accelerated bootstrap interval (BCa interval) method, which is a second-order accurate interval that addresses these problems (Efron 1987; Huang et al. 2007). All the results reported in this paper have been confirmed by bootstrap permutation test (*n* = 200) with significance level of 0.05, which shows consistency with very small variances.

The ROC curves allow us to explore the effects of different pFDR choices on the false positive and false negative rates. ROC curves are usually summarized by the Area Under Curve (AUC). Since we are only interested in low FDRs, we summarize the ROC curves with the partial area under the ROC curve.

The first metric to quantify the estimation performance is the standardized partial AUC (*spAUC*) (McClish 1989), which is independent of the level of the specified false positive rate (FPR). It is defined in the following equation, where *e* is the FPR level we are evaluating:

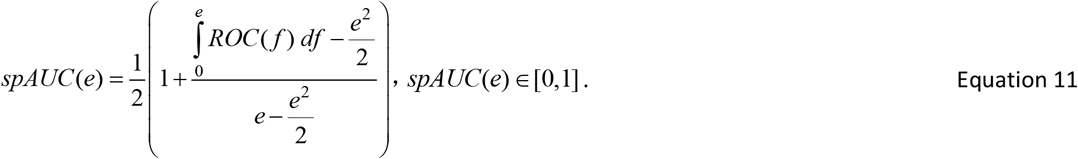

*spAUC* varies from 0 to 1, with 0.5 and 1 are the random and perfect diction. We use *spAUC*(0.1) in latter summary of results.

The second metric is the Performance Improvement Ratio (**PIR**), which compares the *spAUC*(*e*) of one method (M1) against another method (M2) with the same baseline. It is defined as follows:

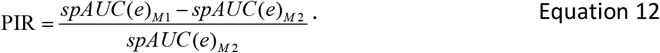

**PIR** can be positive or negative. Positive **PIR** indicates that M1 outperforms M2, and vice versa.

### 5 Source Connectivity Simulation Module: EECoG_scSim

Simulation is useful when we need to select the best available methods in a certain condition. For example, we have 4 ESI methods plus ECoG Laplacian, but we never know the “ground truth connectivity” beneath the real data or which is the most suitable method for our particular data. The only way to answer this question is to create a simulation as similar as possible to the experimental condition where the data is recorded. EECoG_scSim is designed to do so by simulating a structured partial correlation matrix as a simulated true source connectivity, and then generate the source activities with such connectivity. By using the EECoG lead fields generated from EECoG_FWM, we are able to simulate the same experiment with known source connectivity. Together with EECoG_conComp, EECoG-Comp provides tools for both testing methods and exploring data.

EECoG_scSim provides human based simulation for validation as well. A standard human forward model based on the MNI152 template (http://www.bic.mni.mcgill.ca) and standard bioSemi electrodes layouts is calculated with Boundary Element Method (BEM) from BrainStorm (Tadel et al. 2011), and the corresponding lead field is provided for human simulation. EECoG_scSim allows the users to change the standard EEG electrode layouts from 10-20 system to 256 electrodes of full head coverage as well as the size of the cortical grid.

The theory behind this simulation is the same as the forward modeling procedure. The simulated sensor time series ***V**_LF_* for a given head model ***LF*** is generated from:

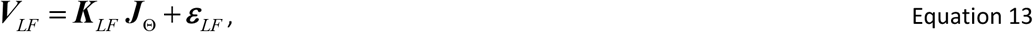

where ***J**_Θ_* is the current source density sampled independently from distribution *N* (***0***, *Θ*^−1^), with *Θ* being the underlying PCM. ***K***_*LF*_ is the lead field for a given head model. *ε*_*LF*_ is the sensor noise with *ε*_*LF*_ ~ *N*(0, ξ × ***I***). The measurement signal to noise ratio (SNR) is 10db for EEG and 20db for ECoG. The length of each cortical grid time series can be the same as the real recordings.

Simulations of ***J***_Θ_ are based on two different structured PCMs:

1. Sparse random PCM Θ_*rand*_, where a small number of the sources on the cortical grids are connected randomly;
2. Sparse block diagonal PCM Θ_*block*_, where a small number of patches of cortical grid points are active and densely connected between them.

The lead fields used in the simulation module are ***K**_H, eeg_* (Human EEG), ***K** _M, eeg_* (Monkey EEG), ***K** _M, ecog_* (Monkey ECoG) and ***K** _Lap_* (Monkey ECoG Laplacian “pseudo lead field”).

### 6 Specification of Simulations for the MDR dataset

We first evaluate the performance of all these methods in EECoG_scSim for both standard human EEG and monkey EECoG by means of simulations (with different number of sources, sensors and partial correlation matrix (PCM) structures); Then, we choose the best method from the monkey EECoG simulation to compared against; At last, we use all these ESI methods in exploring the source connectivity of the real data and give the final ranking of all these tested ESI methods as results.

These simulations are started with the generation of structured PCMs of the neural sources (or generators on the cortical mesh) denoted as *Θ*, and they can have random and block structure which is specified as Θ_*rand*_ and *Θ*_*block*_ (more descriptions in section 2.5). For simplicity, all symbols without specific suffices refer to general definition rather than a specific case—suffixes will be added only when necessary. Based on these two types of PCMs, the source activities ***J*** can be generated as 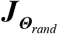 and 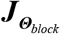 without source noise. Both *Θ* and ***J*** are the same for all human and monkey simulations as the simulated “truth”. The different simulation settings are distinguished by the first term of suffices, *HS* for Human Simulation and *MS* for Monkey Simulation. We latter use *MD* to refer to real Monkey Data analysis. The sensor signals ***V*** are generated by multiplying the corresponding lead field ***K*** with the source activities ***J*** (corresponding to a given PCM *Θ*) with sensor noises ε added, this procedure can be described with the following equations:

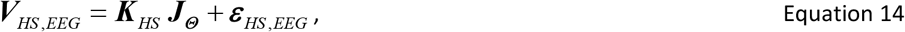

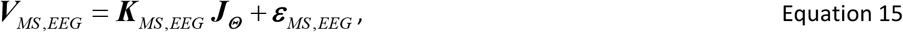

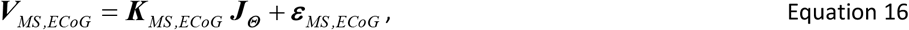

***V**_HS, EEG_* is the simulated human EEG with dimension *N*_*HS, chan*_ × *N*_*tSample*_, *N_HS, chan_* ε (19, 32, 64,128) is the number of human EEG channels, and *N*_*tSample*_ = 10800 is the number of time samples. The sampling frequency is 400Hz for all simulations, which is kept the same as that in the preprocessed monkey recordings. ***K**_HS, EEG_* is the Human EEG lead field with dimension *N*_*HS, chan*_ × *N*_*HS,sour*_, *N*_*HS, chan*_ is the number of human EEG sensors and *N*_*HS,sour*_ ε (150, 200, 300, 350). *ε*_*HS,EEG*_ is the human EEG sensor noise which follows I.I.D. Gaussian distribution *ε*_*HS,EEG*_ ~ *N*(0,ξ × *I*) with dimension of *N*_*HS, chan*_ × *N*_*tSample*_. ***V**_MS, EEG_* and ***V***_*MS, ECoG*_ are generated in the similar way but with monkey EEG lead field ***K***_*MS, EEG*_ (*N_MS,EEG_* × *N_MS, sour_*) and monkey ECoG lead field ***K**_MS, ECoG_* (*N_MS, ECoG_* × *N_MS,sour_*). In order to select the best method to use on the actual monkey EECoG data (***V**_MD,EEG_* and ***V**_M,DECoG_*), we use the same number of the available monkey EEG channels *N*_*MD,EEG*_ = 15 and ECoG channels *N*_*MS, ECoG*_ = 125, as in the simulation, but changing the number of sources *N*_*MS,sour*_ ε (150, 200, 300, 350), note that, *N*_*MS, ECoG*_ < *N*_*MS, Sour*_ and *N*_*MS, EEG*_ < *N*_*MS, Sour*_ always hold.

Then, we apply the 4 well-known Electrophysiological Source Imaging (ESI) methods, namely MNE, eLORETA, LCMV and SSBL in human simulation (EEG), and MNE, eLORETA, LCMV and ECoG Laplacian (this last is viewed as another “pseudo inverse method”) in monkey simulations to reconstruct the source activities. The source activity estimator can be denoted as 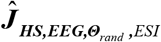 for example, among all the sun-notations: 1) *HS* stands for human simulation, which can also be *MS* for monkey simulation and *MD* for monkey data, this is a notation for experimental condition; 2) *EEG* is the modality, and it can be *ECoG* as well; 3) *Θ*_*rand*_ is the underlying true random source connectivity matrix, which can alternatively be a blocked structured source connectivity matrix *Θ*_*block*_, and it is unknown in the real data analysis; 4) *ESI* stands for different ESI methods, like MNE, eLORETA, LCMV, SSBL and ECoG Laplacian—specified when necessary.

With true source activities in theory ***J*** and estimated 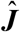, we are able to further estimate the underlying source level connectivity in terms of PCM. We take an example of 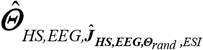 to describe our notations for the connectivity estimators: 1) The first suffix indicates the type of experimental condition, which can be *HS* for Human Simulation, *MS* for Monkey Simulation and *MD* for Monkey Data analysis; 2) The second suffix is modality, which can be EEG or ECoG; 3) The third suffix is the source activities based on which this source connectivity matrix is estimated, for example it can simply be 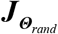, which means the direct estimation from the true source activities with underlying connectivity *Θ*_*rand*_, or like 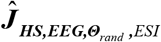 in this example, which is the estimator from the reconstructed source activities obtained with a specific ESI method *ESI* for human EEG simulation (*HS*) with the underlying true source connectivity *Θ*_*rand*_.

We clarify that with this simulation we produce the true source activities ***J_Θ_*** (no volume conduction) and its associated underlying true source connectivity *Θ*. Before further analysis, we bandpass all the true source activities from 8-12Hz to limit the connectivity estimations to alpha band. We then proceed to use the graphical LASSO (Hsieh et al. 2014) to find the “estimated true source connectivity” 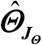 which will help us quantify the error due to the partial correlation estimation procedure, without additional errors due to the source reconstruction.

With this machinery in place, we can generate simulated recordings of human EEG, monkey EEG and monkey ECoG based on models with different number of sources and different number of sensors. After band pass filtering, we apply different ESI methods to determine the “estimated source activities” 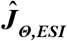 and finally apply the graphical LASSO to estimate the “ESI source connectivity” 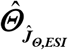.

To summarize the ESI methods performances in simulation, where we know the true source activities and connectivity, we use Performance Improvement Ratio (**PIR**_*ESI*_) for a specific ESI method as a metric. For this case, M1 is the ESI estimated source connectivity and M2 is the estimated true source connectivity. For the analysis of the real MDR data, where we do not have true source activities and connectivity, we only report *spAUC*(0.1)_*ESI*_, which compares the ESI estimated source connectivity against the best ESI source connectivity estimator from the monkey simulations.

All these metrics are compared in the same experimental condition (like human simulation *HS*) with the same modality (like EEG), thus we omit these suffices here. To be specific in this simulation, **PIR**_*ESI*_ is defined as the accuracy improvement ratio of source PCM estimation 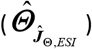 from the reconstructed source activities from a specific ESI 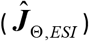 against the direct true source PCM estimation 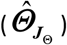 from the true source activities (***J***_*Θ*_). The accuracy is measured by *spAUC*(0.1) (see section 2.4 for more details) and they are written as *spAUC*(0.1)_*ESI*_ and *spAUC*(0.1)_*J*_ respectively, which are calculated for both 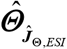 and 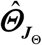 by contrasting them with the true source connectivity matrix (*Θ*). **PIR**_*ESI*_ is calculated as follows:

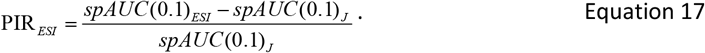

Positive **PIR**_*ESI*_ indicates that this particular *ESI* method outperforms the direct estimation from the source activities, and vice versa. The whole simulation procedure is illustrated in the flowchart below in Fig.6, and the results will be present in next section.

**Figure 6.**
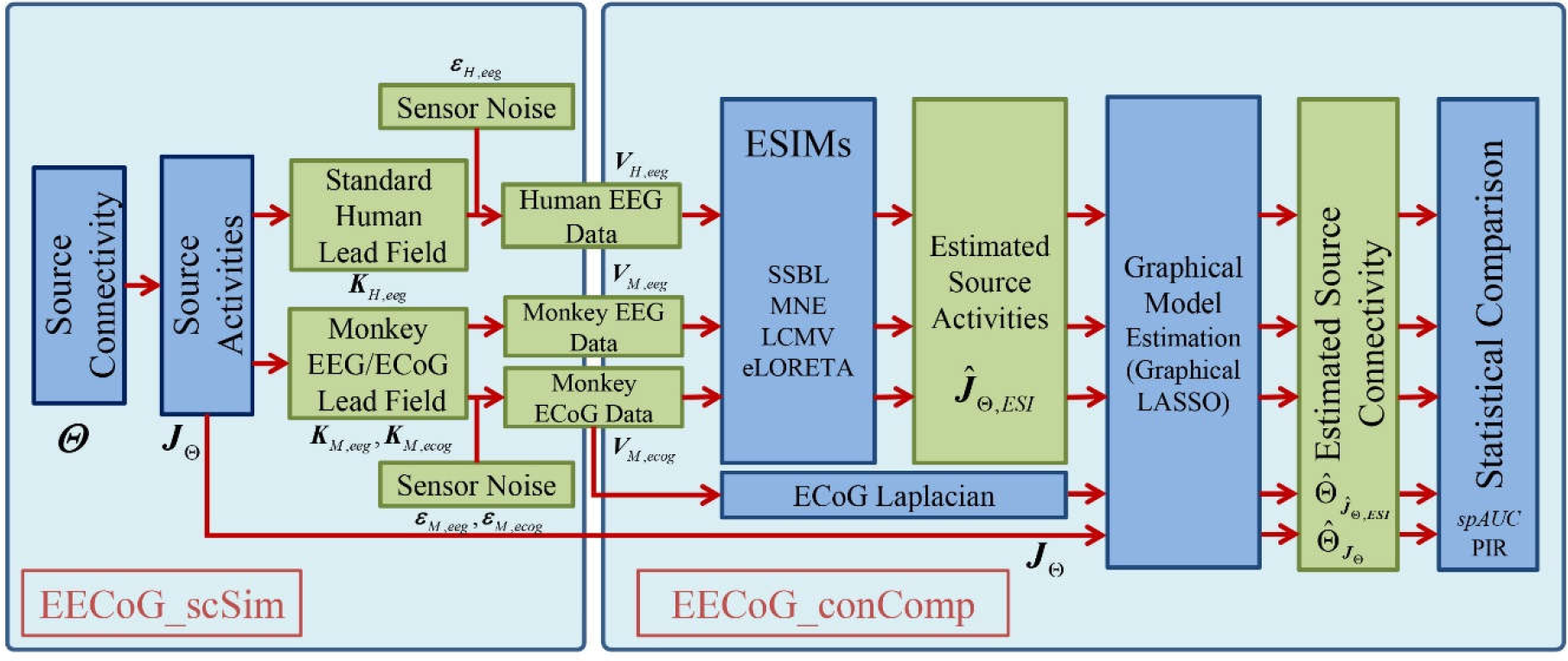
The flowchart of the simulations to compare ESI methods. Firstly, we use EECoG_scSim to generate simulations of human EEG ***V**_H,eeg_* and monkey EECoG ***V**_M, eeg_* and ***V**_M, ecog_* with two types of known structured source connectivity (*Θ*) and true source activities (***J***_*Θ*_); Secondly, we use EECoG_conComp to compare the estimated source connectivity in terms of partial correlation matrix (PCM) 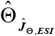 from the estimated source activities 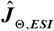 from different ESI methods; Thirdly, compare the connectivity estimators from different ESI methods 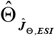 against that 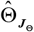 estimated from the simulated true source activity ***J**_Θ_*, and find the method with the highest Performance Improvement Rate **PIR**_*ESI*_, which is ECoG Laplacian; Finally, we compare all the ESI methods against the ECoG Laplacian as a standard to compare against in latter real data validation and report the results in terms of standardized partial Area Under the ROC Curve with False Positive Rate of 0.1 *spAUC*(0.1).

## 3. Results

### Human EEG Simulation

In the human simulation, we are testing the performance of different ESI methods on the standard human head model (MNI152 template http://www.bic.mni.mcgill.ca) with different electrode layouts and different sizes of cortical grids. In this simulation, the number of sensors is *N*_*chan*_ ∈(19, 32, 64,128) for 10-20 system, 32,64, and 128 channel system with whole head coverage, the number of sources is *N*_*sour*_ ∈ (150, 200, 300, 350) over the whole cortex. Due to the computational load of graphical model estimation and ROC evaluation (bootstrapping for ROC confidence band), we are only simulating small cortical grids, therefore we have 16 results in total (4 number of sensors times 4 grid sizes) for each type of source connectivity setting.

The results showed the ESI methods performance improvements in terms of PIR_*ESI*_ for both random and block sources for various sensor and source configurations. In summary, PIR_*ESI*_ of all the ESI methods improved less than 10%. The best performance was always achieved when the source connectivity had a block structure.

### Simultaneous Monkey EECoG Simulation

EECoG-Comp platform allows us to simulate simultaneous monkey EECoG data to test not only the ESI methods but also ECoG Laplacian described in Section 2.4. Due to the fixed number of EECoG channels in the real experiment, we only tested the performance of ESI methods across different sizes of cortical grids: *N*_*sour*_ ∈ (150, 200, 300, 350), the methods tested in this simulation are MNE, LCMV, eLORETA and ECoG Laplacian. This means for each data (EEG or ECoG), we have 4 groups of results for each type of structured source connectivity.

The results from this simulation showed that the ECoG Laplacian almost always had the largest improvement, and the only exceptions were EEG and ECoG with 150 sources. In total, the performance differences between the ESI methods and the Laplacian are within 5%. Though ECoG Laplacian improves the source connectivity estimation the most, the results was not impressive and might not have practical implications. Like the human simulation, the best performance was always achieved when the source connectivity had a block structure.

### ESI methods Comparison with the Real Simultaneous EECoG MDR Data

Even though the ECoG Laplacian connectivity estimation did not yield huge performance improvement in the simulations, it did provide the highest PIR_*ESI*_. Therefore, we choose the ECoG Laplacian as a yardstick (even though we know it is an imperfect “ground truth”) when exploring the real data and compare the ESI methods against ECoG Laplacian.

It is also interesting to see that all the methods are giving “higher *spAUC* for the awake state data, and ECoG outperforms EEG more than 10%. On the contrary, in anesthesia state, all the methods give equally low accuracy compared with ECoG Laplacian, there is even no clear performance difference between EEG and ECoG in the anesthesia state data.

**Figure 7.**
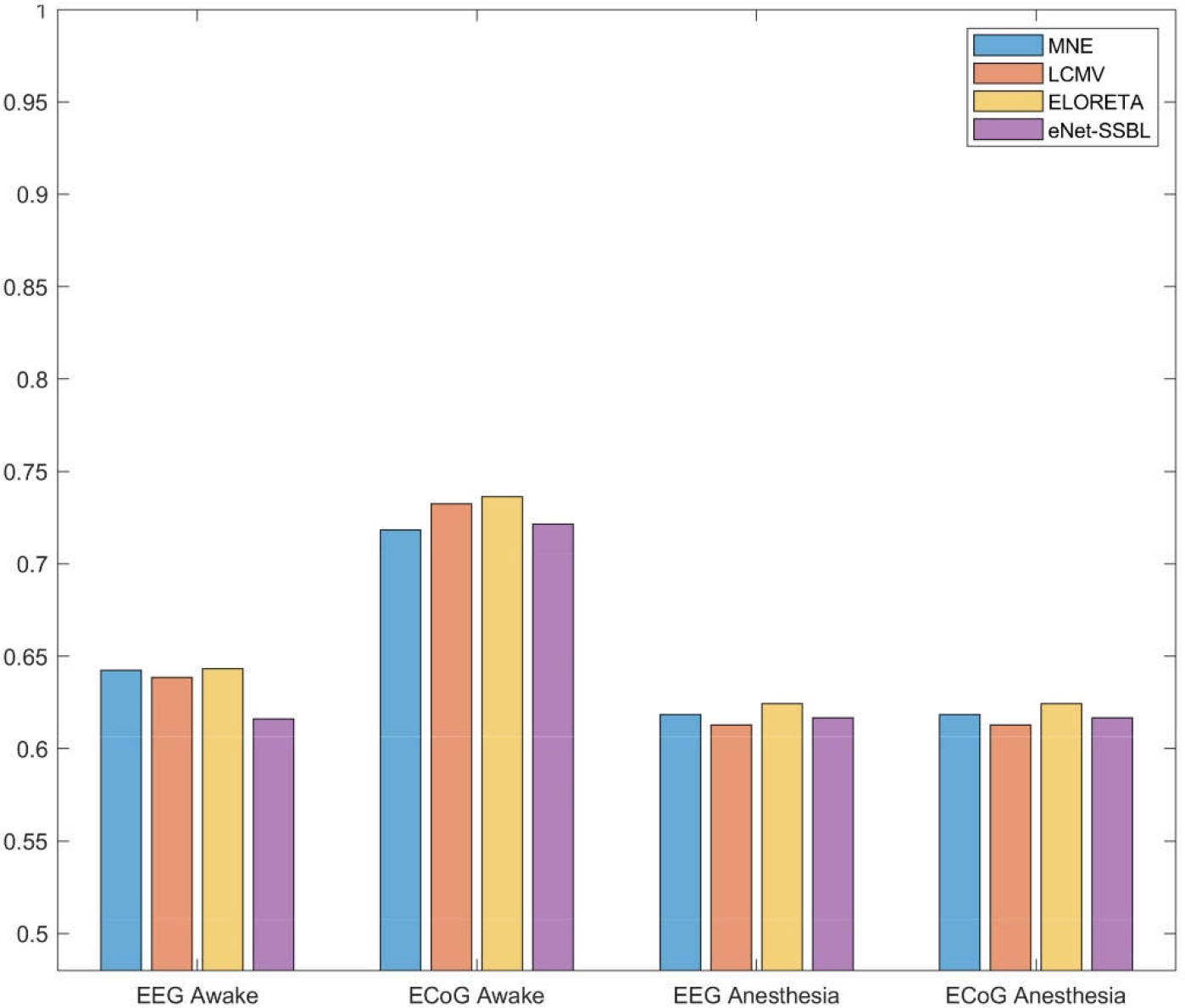
The standardized partial Area Under the ROC Curve with False Positive Rate of 0.1 *spAUC*(0.1)_*ESI*_ of ESI methods performance in simultaneous EECoG MDR data. Each bar stands for one ESI method: MNE (blue), LCMV (orange), eLORETA (yellow) and elastic-net SSBL (purple). Each cluster of bars represents a group of recordings: they are EEG awake, ECoG Awake, EEG Anesthesia and ECoG Anesthesia (from left to right). In the MDR data analysis, we first cleaned the data with EECoG_Preproc, then input the data to EECoG_conComp. By applying different ESI methods, we estimated the source activities and ECoG Laplacian, band passed to alpha band (8-12 Hz), estimated the graphical model parameters with graphical LASSO and do the statistical evaluation of the similarity between the ESI reconstructed partial correlation matrix (PCM) and ECoG Laplacian PCM. The results are summarized in the bar diagram below in Fig. 9 in terms of _*spAUC*(0.1)_. Note that it is a different metric from PIR _*ESI*_, the one we use to compare different ESI methods in. All *spAUC*(0.1) values are within the range of [0.61 ~ 0.74] with mean 0.65 and standard deviation 0.05.

## 4. Discussion

In this paper, we provide EECoG-Comp, an open source EEG and ECoG (EECoG) analysis platform. We illustrate its use with the only publicly available simultaneous EECoG recordings, the MDR dataset (Nagasaka et al. 2011). To our knowledge, this is the only public dataset that combines large coverage ECoG with moderate coverage EEG and high-quality structural imaging data, thus allowing the computation of the lead fields required for evaluation of Electrophysiological Source Imaging (ESI).

We use the EECoG-Comp platform and the MDR dataset to compare source connectivity estimation (with partial correlation) based upon 4 popular ESI methods (MNE, eLORETA, LCMV and SSBL). The comparisons were carried out in several scenarios: a) Using standard human lead fields and neural (source) activity simulated with known partial correlation patterns; b) Using the lead fields from the monkey MDR EECoG data and simulated neural (source) activity as in a); c) Using both the MDR lead fields and actual recordings. It should be noted that we achieved equivalent statistical significance thresholds level across all comparisons. From these comparisons several preliminary conclusions emerge:

1. For real MDR data, 4 popular ESI methods (MNE, LCMV, eLORETA and SSBL) showed significant but moderate concordance with a usual standard, the ECoG Laplacian (standard partial *AUC* = 0.65 ± 0.05);
2. The results from both simulation and empirical data analysis agree with several other papers (Mahjoory et al. 2017; Palva et al. 2018) in which most ESI connectivity estimators suffer from very high level of false positive and false negative connections when compared to the true source connectivity.
3. In simulations, sparse random source connectivity estimation is shown to be very challenging. None of the ESI methods or ECoG Laplacian achieve high levels of accuracy of source connectivity estimation. By contrast, there is a slight improvement in performance for source connectivity estimation that have a “block” type of activations and partial correlations, which may be less sensitive to the effects of “leakage” (Paz-Linares et al. 2018). This is consistent with the results described by Todaro (2018) which quantified the effect of leakage for this same preparation.
4. In simulations, the ECoG Laplacian (Nunez and Srinivasan 2006), which does not depend on the solution of the forward modeling, is only marginally better in performance than any of the 4 ESI connectivity estimators. But it still does not provide high accuracy when compared to the true source connectivity. This result points to the need for both better forward models (Piastra et al. 2018) as well as better methods to deal specifically with accuracy of source connectivity estimations (Palva et al. 2018; Gonzalez-Moreira et al. 2018).
5. We suggest that the ECoG might not be the best tool to investigate the actual primary current density, that future experiments might require other types of measurements like Local Field Potential (Bimbi et al. 2018). Current technological advances may enable this type of wide coverage joint EEG-LFP experiment soon.

While analyzing the data, we need to stress that due to the complexity of the simultaneous EEG and ECoG experimental setup, the data quality is not that ideal. It is striking that careful artifact control and signal processing procedures failed to show a spectral peak in the alpha range (which is commonly observed in monkey experiments such as described in van Kerkoerle et al. (2014), Haegens et al. (2015) and Bollimunta et al. (2011)). This may be due to the circumstances of the recording. The monkey studied was blindfolded but may have been very alert due to unrolled factors. It also could have been due to the peculiarities of this monkey, which sometimes, as in some humans, alpha activity is not present (Charles Schroeder, personal communication). Also, as pointed out by one of the reviewers, and mentioned in the introduction, there is only one data set included in the analyses presented. Therefore, general neurobiological principles cannot be generalized from the results obtained. In addition, the relative merits of one ESI over the other are not conclusive. The small differences observed might not in practice indicate the superiority of one ESI technique over another.

More simultaneous EEG and intracranial experiments like the original MDR are still needed. The design of such experiments can be improved based on our results. For example:

1. We have shown that the EEG channels that have to be rejected are all close to the ECoG connector. Only 15 EEG channels were then left, which is very challenging for inverse estimations, thus we may suggest a higher density recording system (at least 32 channel EEG).
2. The subject is a male monkey, who has very thick muscles on both sides of head (see Fig. 5), which not only makes the forward modeling more difficult (may need to model the muscle as another independent counterpart with different electrical properties) but also introduces more artifacts, thus we may suggest to choose a female subject for the similar experiments.

Thus, the analysis of the MDR data set with this EECoG-Comp platform cannot only serve to compare different ESI methods in both simulations and real data, but also provide suggestions for future experiments. We share this platform via https://github.com/Vincent-wq/EECoG-Comp so that ESI researchers can access, extend and customize it for their own EECoG analysis, and help the neuroscientists to interpreted their data better. As indicated before, for valid neurobiological conclusions future experiments should be carried out with improved experimental design as well as a larger sample of monkeys. It is impossible to obtain general physiological conclusions with such a limited dataset. Nevertheless, the MDR dataset was useful to test our software platform. We certainly hope that the present work will stimulate further high-density concurrent EEG-intracranial recordings in monkey preparations.

## 5. Conclusion

In this paper, we first provide the open source EECoG-Comp platform, which aims to help researchers analyze the source level connectivity of simultaneous EEG and ECoG experimental data, to achieve this goal, this platform offers four modules: 1) EECoG_Preproc to clean the recordings and reject artifacts, 2)EECoG_FWM to build the realistic biophysical forward model for EEG and ECoG, 3) EECoG_conComp for inverse solutions, connectivity estimation, and statistical comparison, and 4) EECoG_scSim for source level connectivity simulation and server as the null model for self-validation and exploring the real data. This platform has great potential in solving the following problems: 1) evaluate the effectiveness of EEG analytical methods (compare to ECoG), 2) evaluate different ESI methods (the example we use to demonstrate the usage of this platform), 3) evaluate different connectivity estimators. In order to demonstrate the application of this platform, we take the simultaneous EEG/ECoG data of a single *macaca fuscata* from neurotycho.org as an example, build the head model with FEM, clean the data, and test different ESI methods in both simulations and real data.

The results from the demonstration show the capability of our EECoG-Comp platform in analyzing the such simultaneous EECoG data, and a striking result is that none of the tested ESI methods is able to give accurate source connectivity reconstructions in terms of source partial correlation matrix. This work provides practical suggestions for future protocols. In summary, this platform might help the researchers to explore this type of simultaneous EEG and ECoG data and obtain more reliable and confident conclusions. It may well stimulate further concurrent EEG-ECoG experiments in macaques.

## 6. Open Source and Data Sharing

EECoG-Comp platform is designed with the idea of open source and open data sharing (Poline et al. 2012) and is shared via https://github.com/Vincent-wq/EECoG-Comp, all the results in exploring the MDR data set will be shared upon request, including the MRI images, the segmentation files, the head model files, the lead fields and so on, more details are listed in Table 2.

## Acknowledgements and Author Contributions

The authors would like to thank the helpful discussions with Carsten Wolters, Marios Antonakakis, Maria Carla Piastra, and Laura Marzetti in forward modeling, the discussions with Prof. Peng Ren, Mayrim Vega Hernandez, and Eduardo Martinez Montes with simulation and inverse solutions, and thanks Shiang Hu with the help in reference selection and spectrum decomposition. The authors also would like to thank for the support from the Tri-national axis in normal and pathological cognitive aging project with No. 81861128001, and the funds from National Nature and Science Foundation of China (NSFC) with funding No. 61871105, 61673090, and 81330032, and CNS Program of UESTC with No. Y03111023901014005.

## Appendix

## A1: Mathematical Notations

**Table 3.**
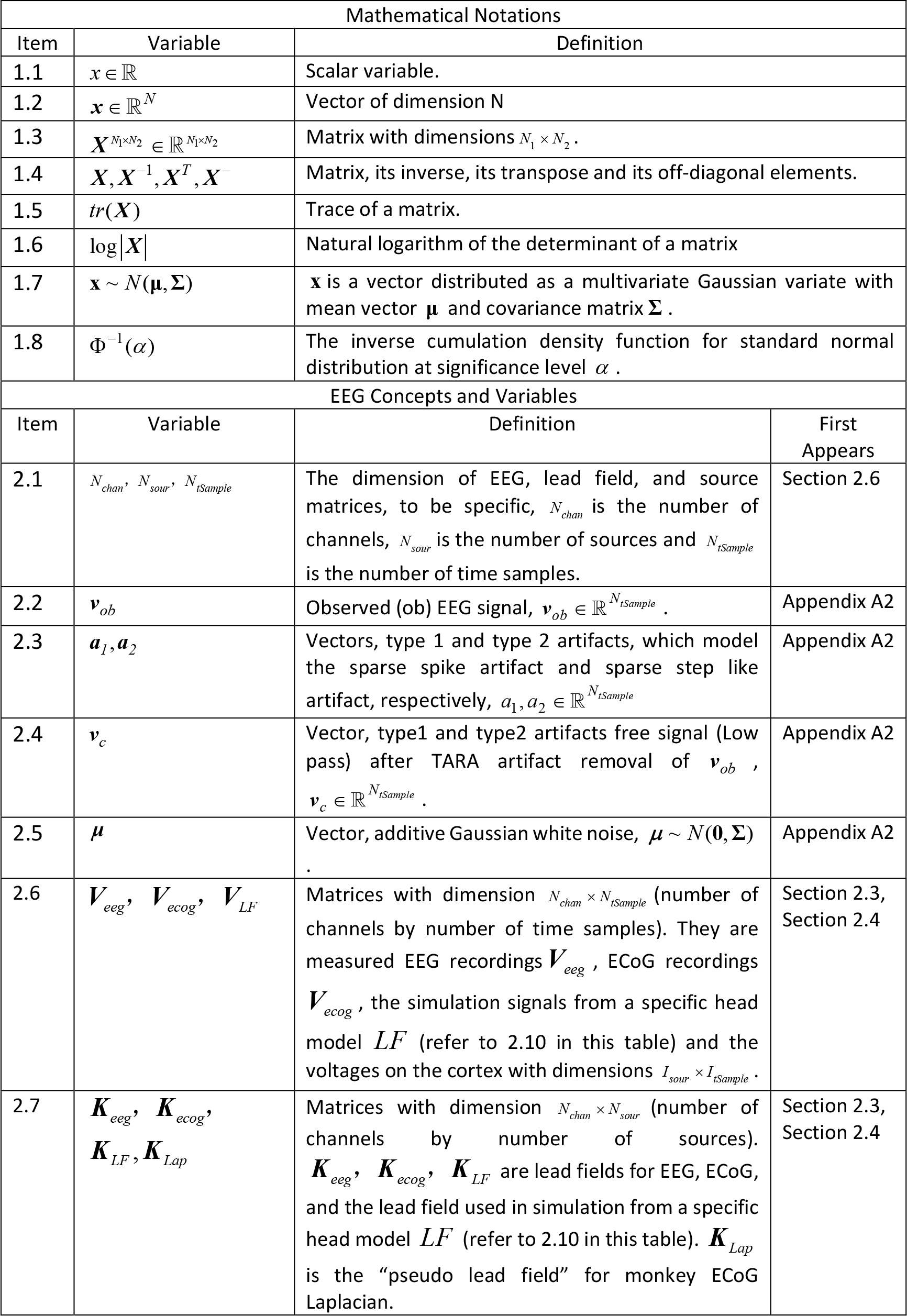
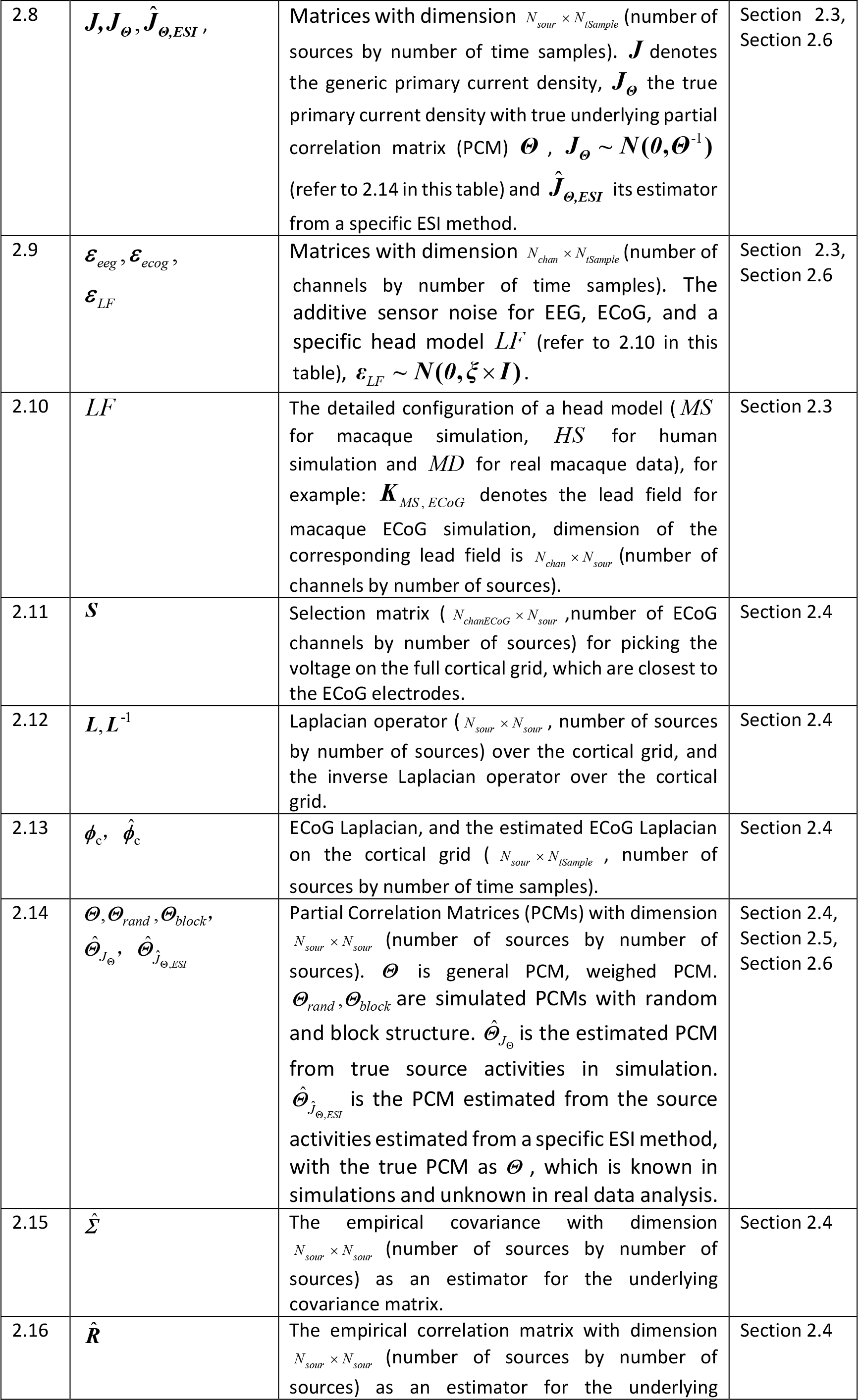
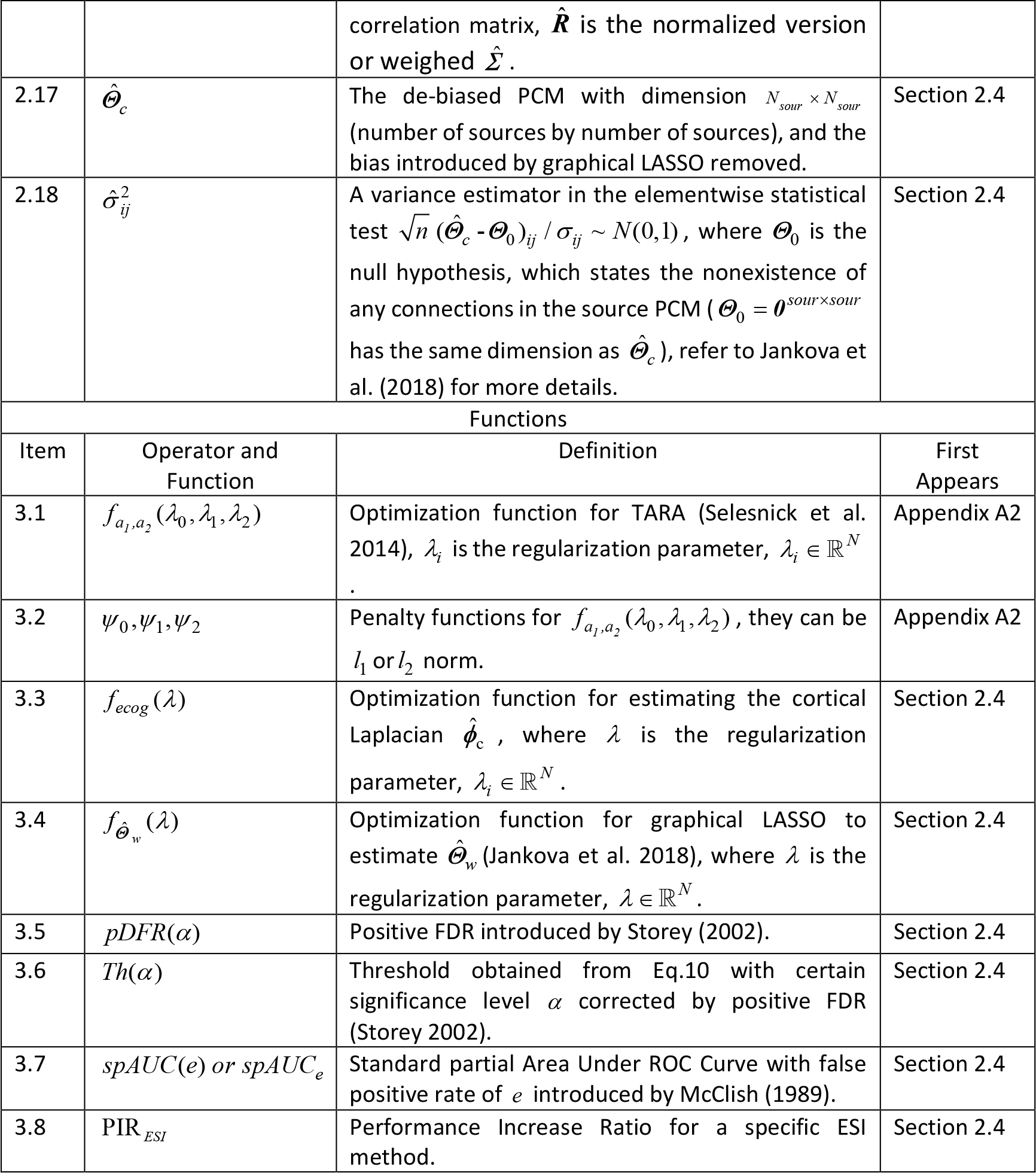
Table of Notations for variables and functions.

## A2: Theory of Transient Artifact Reduction Algorithm (TARA)

We give further brief introduction to the theory behind TARA (Selesnick et al. 2014) in this part. The two types of artifacts that TARA is aiming to remove are: sparse spikes (sparse functions) and step-like discontinuities (functions with sparse first derivatives), and the model (for one channel) can be written as:

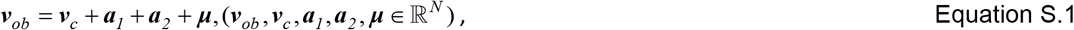

Where ***v**_ob_* is the recorded signal, ***v**_c_* is the low-pass discrete-time signal (artifact free or clean signal) we aim to obtain. ***a**_1_* represents the “spare” spike artifacts (type 1), a small number of spikes compared to the length of signal with also small number of nonzero elements in its first derivative. ***a**_2_* represents the step-like discontinuities (type 2), which also implies a sparse derivative. The term ***μ*** is the additive Gaussian noise. *D* is the first order differential operator. Based on this, the artifact removal procedure can be formulated into an optimization problem:

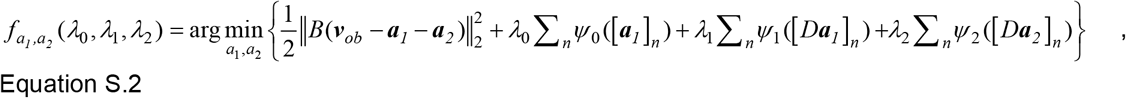

Where *λ*_i_ are the regularization parameters, and ψ_i_ is the penalty function promoting sparsity (or smoothness), which can be ***l**_1_* or ***l**_2_* norms (depend on the characteristic of the real data and the assumption of the artifact free signal ***v**_c_*). *B* denotes the high-pass filter annihilating the low-pass (artifact free) signal ***v**_c_*, and it can be estimated as:

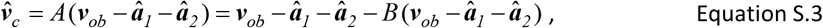

where *A* is defined as the low-pass filter: *A* = ***I*** - *B*.

## Footnotes

1 Qing Wang and Pedro Antonio Valdés-Hernández are co-first authors, who contributed equally to this paper.

2 Noteworthily, more recent and realistic phantoms, even 3D printed, are now available for these tests (Collier et al. 2012).

